# Cryo2StructData: A Large Labeled Cryo-EM Density Map Dataset for AI-based Modeling of Protein Structures

**DOI:** 10.1101/2023.06.14.545024

**Authors:** Nabin Giri, Liguo Wang, Jianlin Cheng

**Author notes:** Corresponding author: Jianlin Cheng.

## Abstract

The advent of single-particle cryo-electron microscopy (cryo-EM) has brought forth a new era of structural biology, enabling the routine determination of large biological molecules and their complexes at atomic resolution. The high-resolution structures of biological macromolecules and their complexes significantly expedite biomedical research and drug discovery. However, automatically and accurately building atomic models from high-resolution cryo-EM density maps is still time-consuming and challenging when template-based models are unavailable. Artificial intelligence (AI) methods such as deep learning trained on limited amount of labeled cryo-EM density maps generate inaccurate atomic models. To address this issue, we created a dataset called Cryo2StructData consisting of 7,600 preprocessed cryo-EM density maps whose voxels are labelled according to their corresponding known atomic structures for training and testing AI methods to build atomic models from cryo-EM density maps. It is larger and of higher quality than any existing, publicly available dataset. We trained and tested deep learning models on Cryo2StructData to make sure it is ready for the large-scale development of AI methods for building atomic models from cryo-EM density maps.

## Background & Summary

Accurately determining three-dimensional (3D) structure of proteins is critical for unlocking key insights into their molecular functions^1^ and interactions with other proteins as well as small molecules like ions, ligands, and therapeutic drugs^2^. In recent years, cryo-EM^3^ has emerged as the most important technology for experimentally determining the structures of large protein complexes that are difficult or impossible for other experimental techniques such as X-ray crystallography or Nuclear Magnetic Resonance (NMR) to solve. The field of cryo-EM is advancing at a rapid pace with improvements in image data collection and processing techniques^4^. This progress has resulted in large amount of high-quality cryo-EM images of proteins and their complexes, and high-resolution 3D density maps reconstructed from two-dimensional (2D) images. EMPIAR, the Electron Microscopy Public Image Archive^5^, is an archive for raw images, such as micrographs and particle stacks^6^, that is used in the construction of cryo-EM density maps. Subsequently, these reconstructed 3D cryo-EM density maps are deposited into EMDB, the Electron Microscopy Data Bank.^7^.

However, the building of atomic 3D models of large protein complexes from cryo-EM density maps is still a very challenging problem that often requires extensive manual intervention. The task is particularly difficult if the structures of the individual chains (units) in large protein complexes are not available or cannot be predicted from sequences by cutting-edge protein structure prediction methods such as AlphaFold2^8^ which serve as templates to fit into cryo-EM density maps for building the models of the complexes.

To address this problem, significant efforts have been put into developing machine learning, particularly deep learning (DL) methods^9^, to directly build atomic models from cryo-EM density maps as the density in each voxel (i.e., 3D pixel) of the high-resolution density map contains sufficient information about protein structures in most cases. Therefore, sophisticated deep learning models trained with density maps as input and their corresponding structures as labels can potentially infer protein models that fit the cryo-EM density maps correctly and accurately^10^.

However, designing and implementing deep learning-based methods for building atomic models from cryo-EM density maps requires a large labeled cryo-EM dataset. Different methods have been trained previously to predict different aspects of protein structures from cryo-EM density maps, nevertheless, up to now, there is still no work on generating a publicly available, large, labeled dataset to push the frontier of the model building from cryo-EM density maps. Recently, the exponential growth of cryo-EM has led to a surge in the deposition of high-resolution cryo-EM density maps in the EMDB^7^ and their corresponding protein models in the Protein Data Bank (PDB)^11^. Leveraging these precious resources, we created Cryo2StructData^12^, a large labeled cryo-EM density map dataset for developing and testing AI-based methods to build atomic models from cryo-EM density maps. We also trained and tested deep learning models on the dataset to rigorously validate its quality. Cryo2StructData is the first, large, publicly available cryo-EM dataset with standardized input features and well-curated output labels that is fully ready for the development of AI-based atomic structure modeling tools in the field.

## Methods

### Related Works

#### Protein Structure Modeling from Simulated Density Maps

Early methods to build atomic models from cryo-EM density maps utilized protein structures in the Protein Data Bank^11^ to generate theoretical density maps at different resolution, usually referred to as simulated density maps for training and testing. For instance, Cascaded-CNN^13^ utilized `pdb2mrc` from the EMAN2 package^14^, and VESPER^15^ utilized `pdb2vol` from the Situs package^16^ to generate simulated density maps. However, simulated density maps lack the complexity of real-world density maps such as high noise, missing density values, and experimental artifacts that arise from protein particle picking^17^ errors, or atom movement during image capturing in the cryo-EM data collection. As such, deep learning models trained on simulated density maps may not perform well on noisy experimental density maps. Therefore, real-world cryo-EM density maps with labels need to be created to further advance the field.

#### Protein Structure Modeling from Experimental Density Maps

As more and more experimentally determined density maps became readily available in the EMDB, different methods have utilized these maps to predict atoms in cryo-EM density maps, as shown in Table 1. A recent method, DeepTracer^18^, offers users the ability to build models through a web interface. It utilized 1,800 experimental maps to predict the positions and amino acid types of C*α* atoms of proteins, which were then aligned with protein sequences to model protein structures. ModelAngelo^19^ used 3,715 experimental maps for training and a test set of 177 maps for testing. Both DeepTracer and ModelAngelo automatically generate the atomic models of the protein, but they have not released their processed data. Haruspex^20^ used 293 experimental maps for training and an independent test set of 122 experimental maps for testing, and EMNUSS^21^ used 120 and 43 experimental density maps for training and testing respectively. CR-I-TASSER^22^, EMNUSS^21^, and Emap2sec^23^ employed a hybrid approach that combined both simulated maps and experimental maps in their training and validation processes. Haruspex, EMNUSS, and Emap2sec are designed for secondary structure prediction, but they have not made their processed data available.

**Table 1.** Summary of the currently used number of experimental cryo-EM density maps in the datasets for various AI-based methods and their public availability. The dataset size reported for Cryo2StructData^12^ corresponds to the version released as of March 2023.

|  | EMNUSS<sup>21</sup> | Haruspex<sup>20</sup> | DeepTracer<sup>18</sup> | ModelAngelo<sup>19</sup> | Cryo2StructData |
| --- | --- | --- | --- | --- | --- |
| Dataset size | 163 | 415 | 1,800 | 3,892 | 7,600 |
| Data released | × | × | × | × | ✓ |

Furthermore, despite the significant progress, the datasets used in these works only account for a small portion of high-resolution density maps available in the EMDB and they are not publicly available for the AI and machine learning community to use. Significantly, Giri et al.^9^ emphasize the critical importance of developing high-quality cryo-EM datasets, specifically for training and evaluating deep learning methods. In this context, we aim to prepare, preprocess, label, validate, and consolidate complementary information that can be utilized for accurate atomic model building through AI-based methods from experimental cryo-EM density maps.

### Cryo2StructData Preparation

The Cryo2StructData was prepared by a data processing pipeline shown in Figure 1. The data generated from each stage of the pipeline was verified to ensure the details of original experimental density maps were preserved. The source code of the data preparation pipeline is released at the GitHub repository of Cryo2StructData for users to reproduce the process. The details of each data preparation stage are described in the following subsections.

**Figure 1.**
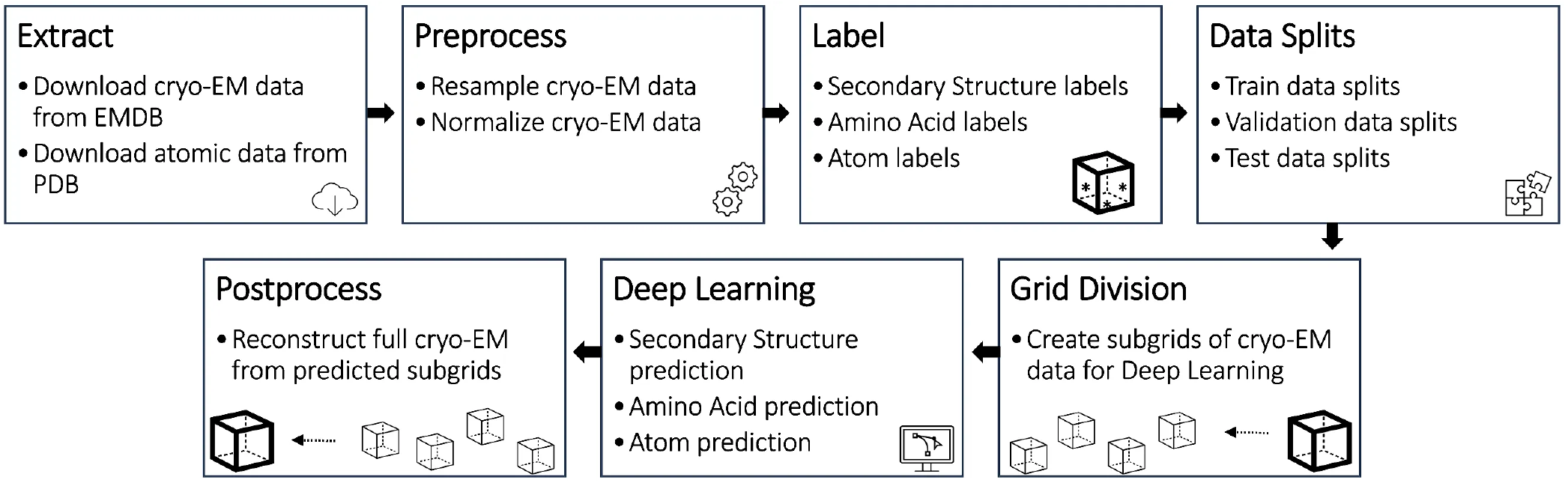
The data preparation and evaluation pipeline for Cryo2StructData.

#### Data Extraction

Cryo2StructData was curated from the experimental cryo-EM density maps released till 27 March 2023. We downloaded relatively high-resolution cryo-EM density maps for proteins and their complexes with resolutions between 1 and 4 Angstrom (Å). In total 9,500 cryo-EM density maps were collected from the EMDB^7^. Similarly, we downloaded the atomic models (i.e., PDB files) corresponding to the density maps from the PDB^11^. The downloaded PDB files were used to label voxels of density maps.

We further filtered out the density maps that do not have structures in the PDB. After the filtering, 7,600 cryo-EM density maps were left for further processing. A supplementary document containing the meta information such as the corresponding PDB entry for each density map, resolution of the density map, structure determination method, software used to determine the density map, the title and the journal of the article describing the density map is provided in the Cryo2StructData Dataverse^12^.

#### Preprocessing

The experimental cryo-EM density maps are stored in MRC^24^ format. The MRC format contains the data in a 3-dimensional (3D) numpy array, often referred to as 3D grid. Each voxel also known as 3D pixel contains a value indicating how strong an atom’s signal is present in the voxel. We refer to these density maps as original raw density maps, which need to be further preprocessed so they can be used to train AI methods. Building upon established practices and techniques^13,18^, the preprocessing is implemented in two steps: (a) cryo-EM density map resampling and (b) cryo-EM density map normalization, as follows:

##### Resample cryo-EM density maps

Different raw cryo-EM density maps usually have different voxel sizes, which need to be standardized. We resampled the density maps to a uniform voxel size of 1 Angstrom (Å), using *vol resample* command within UCSF ChimeraX^25^ in the non-interactive mode. The idea of resampling density map is illustrated in Figure 2a and 2b. All the resampled density maps have uniform, the same voxel size and are used for the following normalization step.

**Figure 2.**
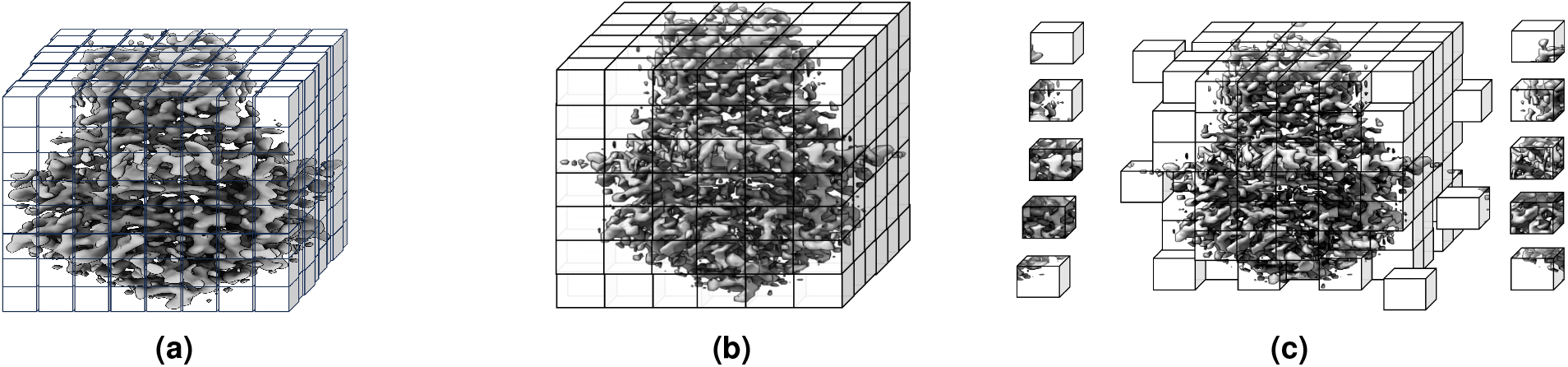
An example of density map grid resampling and division of a cryo-EM density map. (**a**) Density map (EMD-22898) in the original grid. (**b**) Resampled density map to the uniform grid size of 1Å. (**c**) Grid division of the density map where each cube is sub-grid with specific size, in this illustration, it is 1 Å× 1 Å× 1 Å.

##### Normalize cryo-EM density maps

We applied scaling and clipping to normalize the density values of the resampled cryo-EM density maps. In the cryo-EM density maps, positive density values represent regions (voxels) where the protein is likely present. The range of these values can differ from maps to maps with some maps containing values in one range (e.g., [-2.32, 3.91]) and others in another range (e.g., [-0.553, 0.762]). To make the density values of different density maps comparable, we perform the percentile normalization by first calculating 95^*th*^ percentile of the positive density values in a density map and then dividing the density values in the map by this value. To deal with the extreme outliers in the density values, we set all values below 0 to 0 and all values above 1 to 1 after the division. The normalization removes the cross-map difference caused by differences in experimental conditions, and software used to process the maps, allowing AI methods to learn patterns across different cryo-EM density maps. Finally, the resampled and normalized map is saved in the MRC format^26^ with the filename as `emd_normalized_map.mrc`. This file is used as an input to deep-learning model and for label generation process.

#### Labeling Density Maps

Each voxel of a cryo-EM density map has a density value which positively correlates with the possibility of the presence of an atom’s signal in the voxel, which can be used as input features to predict positions and types of atoms. After a broad review^9^ of the structural properties of current machine learning methods to build atomic models from density maps, we created the following labels for voxels in a density map: atom labels, amino acid labels, and secondary structure labels.

For atom labeling, we created an empty (with values of 0) mask with the same dimension as the `emd_normalized_map.mrc`, indicating that the mask was empty. We then utilized the corresponding PDB file for the density map to label each voxel containing backbone carbon-*α* atom (C*α*) as 1, backbone nitrogen atom (N) as 2, and the carbonyl backbone carbon atom (C) as 3. The mask created from the `emd_normalized_map.mrc` map is a 3D grid, where the location of each voxel is determined by indices (*i, j, k*). But the corresponding PDB file used to label the voxels are in another 3D coordinate system (*x, y, z*). Therefore, we calculated the corresponding indices of each backbone atom (C*α*, N and C) in the mask from it’s atomic coordinates using the Formula 1, where *i, j, k* are the grid indices of the atom in the mask, *x, y, z* are the coordinates in the PDB file, *origin*_*x*_, *origin*_*y*_, *origin*_*z*_ are the origin of *x, y, z* axis respectively found in the `emd_normalized_map.mrc` map, and *voxel*_*x*_, *voxel*_*y*_, *voxel*_*z*_ are the voxel size of *x, y, z* axis respectively found in `emd_normalized_map.mrc` map.

Similar to the atoms labeling, we created a second mask containing amino acid type labels. We labeled each voxel containing a C*α* atom with one of 20 different types of standard amino acids, represented by 20 numbers from 1 to 20, while 0 denotes the absence of C*α* atom or unknown amino acid type. Moreover, following the same approach used for atoms and amino acid types labeling, we created a third mask for secondary structure labels. We used UCSF ChimeraX^25^ to identify and extract the secondary structure type (coil, *β* -strand, and *α*-helix) for each C*α* atom from the PDB file. The extracted secondary structures were then used to label each voxel of the mask that contains a C*α* atom. We used 1, 2, and 3 to represent coil, *α*-helix, and *β* -strand, respectively. Figure 3 shows an example of density map labels.

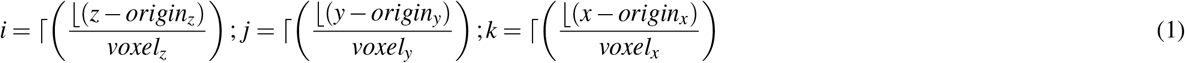

**Figure 3.**
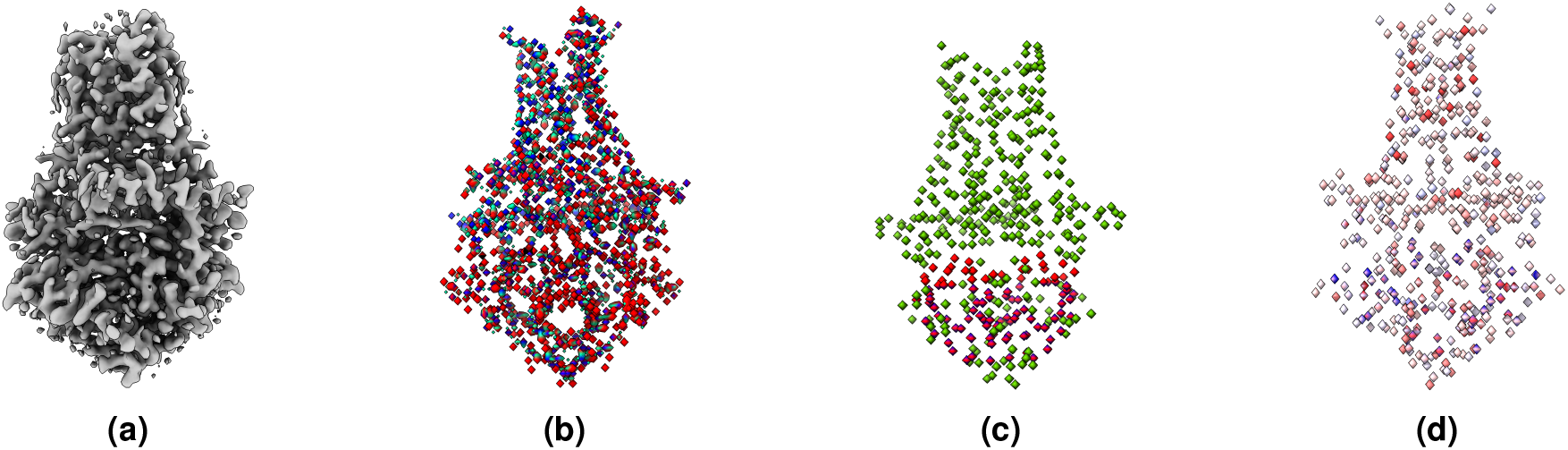
An example of labeling a cryo-EM density map. (**a**) The density map of SARS-CoV spike gycoprotein, EMD-22898 visualized at recommended contour level of 0.7. (**b**) Three different types of protein backbone atoms (C*α*, N, C) labeled in different colors. (**c**) Three different secondary structure elements labeled in different colors. (**d**) Twenty different amino acid labeled in different colors. The image are generated using UCSF ChimeraX’s^25^ surface color by volume data value.

#### Data Splits

We selected 208 cryo-EM density maps, which had previously been used as test data in the previous works^18,27^, from our dataset to create the test dataset^28^. The remaining 7,392 density maps were split into the training dataset consisting of 6,652 maps and the validation dataset consisting of 740 maps according to 90% and 10% ratio. Table 2 reports the number / percentage of maps in the training and validation datsets for each resolution range, indicating the statistics for the two datasets is largely consistent. The resolution value of a cryo-EM maps refers to the level of detail at which the structural features of the biological molecules can be visualized within the 3D map. A smaller resolution value indicates higher resolution and better definition of structural features, while a larger resolution value corresponds to lower resolution and less distinct features. As such, we used resolution-based splitting of the cryo-EM density maps for training and validation of our deep learning model. Additionally, the ID-based splits for the full dataset are provided in the Cryo2StructData Dataverse^12^ because the unique IDs assigned to each cryo-EM density map can be influenced by factors such as the type of entry, metadata, and curation criteria. If necessary, users may choose to split Cryo2StructData into training, validation and test datasets differently.

**Table 2.** The resolution distribution of the maps in the train and validation datasets. ‘Full’ denotes the entire dataset^12^, while ‘Small’ denotes a smaller subset of the dataset^29^. The IDs for the training and validation data are provided in the Cryo2StructData Dataverse^30,31^.

| Resolution Range (Å) | Train Set |  | Validation Set |  |
| --- | --- | --- | --- | --- |
|  | Full | Small | Full | Small |
| 1.0 - 2.0 | 62 (0.93%) | 34 (2.02%) | 5 (0.67%) | 4 (2.14%) |
| 2.0 - 3.0 | 2147 (32.27%) | 535 (31.8%) | 222 (30%) | 55 (29.41%) |
| 3.0 - 4.0 | 4443 (66.79%) | 1111 (66.13%) | 513 (69.32%) | 128 (68.45%) |
| Total | 6652 (100%) | 1680 (100%) | 740 (100%) | 187 (100%) |

#### Grid Division

The dimensions of density maps of different proteins vary and are usually too large to fit into the memory of a standard GPU for training deep learning models. Similar to the approach employed in DeepTracer^18^, Haruspex^20^, CR-I-TASSER^22^, Emap2sec^23^, and Cascaded-CNN^13^, we performed grid division to divide the density maps into 3D subgrids with dimension of 32 × 32 × 32 overlapped by 6 voxels on each face of the subgrid to train deep learning methods. We choose the dimension 32 as it is big enough to capture the patterns (e.g. six turns of alpha helix) in the data and is small enough to be used with GPUs effectively. In the inference stage, the predictions for the sub-grids can be stitched back to obtain the prediction for the full density map by concatenating the predictions for the central 20 × 20 × 20 core voxels of the sub-grids. This approach allowed us to preserve the spatial information of the density maps and overcome the abnormal predictions for the voxels being cut off at boundaries of sub-grids during the grid division. Figure 2c shows an example of grid division. The sub-grids of each density map along with their corresponding labeled mask maps are saved as a single entity in numpy array for deep learning training and test. In total, there are ∼ 39 million training sub-grids and ∼ 4 million validation sub-grids. Moreover, the scripts of dividing density maps are provided at the Cryo2StructData’s GitHub repository for users to create sub-grids of their defined dimensions if necessary.

## Data Records

Cryo2StructData Dataverse^12,29^ contains the necessary metadata, 3D density map, and mask files to enable AI experts without much domain knowledge to develop AI methods for modeling protein structures from cryo-EM density maps.

The following data files for each cryo-EM density map are provided in its individual directory:

- The original cryo-EM density map with its corresponding protein model in the PDB format, the original protein sequence in the FASTA format^32^, and the sequence extracted from the PDB file.
- The Clustal Omega^33^ alignment between the original sequence and the sequence extracted from the PDB file. The two sequences are usually highly similar but not identical because some residues in the original sequence may be disordered and has not been modeled (i.e, no x, y, z coordinates) in the PDB file.
- The resampled and normalized cryo-EM density map generated from the original density map. The values of the voxels in this density map were normalized into the range [0, 1] from their values in the original density map and are ready for being used as input for AI models.
- The labeled masks in which the voxels containing the key backbone atoms (carbon-alpha (C*α*), nitrogen (N) and carbonyl backbone carbon (C)), the C*α* atoms only, the twenty different types of amino acids that C*α* atoms belong to, and the three different types of secondary structures (helix, strand, coil) of the C*α* atoms are labeled. The labeled masks can be used as targeted outputs for AI and machine learning models to predict from the resampled and normalized density maps (input).

A comprehensive documentation of the Cryo2StructData is available within the Cryo2StructData Dataverse^12,29–31^. This documentation provides in-depth insights into the dataset, elucidating the composition of data files, the structure of directories, and the overall organization of the dataset.

## Technical Validation

We validated every step of the data processing pipeline as shown in Figure 1 and described in `Cryo2StructData Preparation` section. The source code used to validate data preparation pipeline is released at the GitHub repository of Cryo2StructData for users to reproduce the process.

### Validation of Preprocessing Step

The preprocessing phase of Cryo2StructData preparation performs two steps : (a) cryo-EM density map resampling and (b) cryo-EM density map normalization. We verified both steps for all the cryo-EM density maps present in Cryo2StructData. Additionally, since each cryo-EM density map contains a unique EMDB-ID, we verified the availability of duplicate density maps and found none.

#### Resampling validation

During the initial stage of data preparation, the density maps were resampled to a uniform voxel grid with a voxel size of 1 Å. To assess the accuracy of this resampling process, we performed validation checks to confirm the voxel size of all density maps within the Cryo2StructData. The validation procedure utilized the `mrcfile` package^26^, a Python implementation of the MRC2014 file format^24^, widely utilized in structural biology for storing cryo-EM density maps. The results of the validation phase clearly demonstrated that all cryo-EM density maps in the Cryo2StructData exhibited a voxel size of 1 Å. This validation, in turn, confirms the precision and consistency of the resampling step, substantiating the attainment of the desired voxel size of 1 Å.

#### Normalization validation

To assess the validity of the normalization step in the data preparation process, we performed checks to identify any outliers and ensure that all density values were confined within the range of [0, 1] for each density map. The validation check did not find any outliers, and all density values were found to lie within the specified range [0, 1]. This validation further corroborates the consistent implementation of the preprocessing step, as all density maps in the Cryo2StructData exhibited values within the defined range [0, 1].

Figure 13 displays the statistical characteristics of a density map, both before and after the preprocessing step, as visualized using UCSF Chimera^25^. By analyzing Figure 13, we can further confirm that the voxel size is 1 Å, and the density values fall within the range of [0, 1], as expected. This provides additional evidence of the correct implementation of the resampling and normalization steps.

### Validation of Labeling Step

We validated the labeling step by leveraging the mask map of labeled C*α* atoms and their corresponding known atomic backbone structure. To obtain approximate 3D coordinates (*x, y, z*) from their respective indices (*i, j, k*), we applied Formula 2. The grid indices, (*i, j, k*), were assigned values of either 0 (indicating the absence of a C*α* atom) or 1 (indicating the presence of a C*α* atom) by the data preparation pipeline. After converting the indices to their associated coordinates for each C*α* atom, we observed, as expected, that the protein backbone structure was not aligned. Consequently, a direct structure-to-structure comparison for evaluation was not feasible. To assess the accuracy of the conversion step, we performed a comprehensive search for the converted coordinates within the known C*α* backbone structure. During the search process, we kept track of the number of precise C*α* atoms matching with the known backbone structure. We randomly selected approximately 4,450 (>55%) of the Cryo2StructData to verify the conversion process. We found that around 4,185 (94%) of the maps had exact matching C*α* positions for all the voxels. For the remaining 6% of the maps, more than 93% of the voxels had exact match, while less than 7% of the voxels deviated slightly from the exact match. The loss of precision in a small portion of C*α* positions in a small portion of density maps is expected due to the interchange between floating-point numbers and integers during the labeling and evaluation steps. Therefore, this validation confirm the accuracy of the labeling step, as smaller than 0.5 angstrom (Å) deviations of a small portion of C*α* (about 6% × 7%) positions is expected.

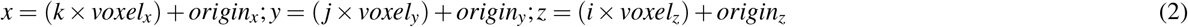

### Validation of MRC files

We conducted a comprehensive validation of the entire Cryo2StructData by executing a series of tests to verify its compliance with the MRC2014 format specification^24^. These tests were performed on the data listed below, which pertains to each density map within the Cryo2StructData:

1. `emd_normalized_map.mrc` : This file is used as an input to the deep learning-based model for training, validation and inference step. As such, this is an important file that needs to pass all the tests.
2. `atom_ca_emd_normalized_map.mrc` : This is the labeled C*α*-only mask map used as labels for training the deep learning-based model, which identifies the C*α* atoms in the input density map.
3. `atom_emd_normalized_map.mrc` : This is the labeled backbone atom (C*α*, C, and N) mask map used as labels for training the deep learning-based model, which identifies the backbone atoms in the input density map.
4. `amino_emd_normalized_map.mrc` : This is the labeled amino acid type mask map used as labels for training the deep learning-based model, which identifies the amino acid residue types of C*α* atoms in the input density map.
5. `sec_struc_emd_normalized_map.mrc` : This is the labeled secondary structure (coil, *α*-helix, and *β* -strand) mask map used as labels for training the deep learning-based model, which identifies the secondary structure types of C*α* atoms in the input density map.

Each of the above files went through a series of tests and checks to identify any potential problems. These files undergoes validation using the `mrcfile` package, specifically the `mrcfile.validate` function. The `mrcfile` Python library is maintained by the Collaborative Computational Project for Electron cryo-Microscopy (CCP-EM)^34^ and serves for reading, writing, and validating MRC2014 files^24^. The following tests were conducted to validate the Cryo2StructData. We provide a brief description of these tests below and refer users to the MRC2014 paper^24^ for detailed explanations of these tests.

1. `MRC format ID string` : The `map` field in the header should contain “MAP”. The character string ‘MAP’ is used to identify file type.
2. `Machine stamp` : The machine stamp should contain one of `0x44 0x44 0x00 0x00`, `0x44 0x41 0x00 0x00` or `0x11 0x11 0x00 0x00`.
3. `MRC mode` : The `mode` field should be one of the supported mode numbers:
  a. **0** 8-bit signed integer
  b. **1** 16-bit signed integer
  c. **2** 32-bit signed real
  d. **4** transform : complex 32-bit reals
  e. **6** 16-bit unsigned integer
  f. **12** 16-bit float
4. `Map and cell dimensions` : The header fields `nx`, `ny`, `nz`, `mx`, `my`, `mz`, `cella.x`, `cella.y`, and `cella.z` must all be positive numbers.
5. `Axis mapping` : The header fields `mapc`, `mapr`, and `maps` must contain the values 1, 2, and 3 - in any order.
6. `Volume stack dimensions` : If the spacegroup is in the range 401-630, representing a volume stack, the `nz` field should be exactly divisible by `mz` to represent the number of volumes in the stack.
7. `Header labels` : The `nlabl` field should be set to indicate the number of labels in use, and the labels in use should appear first in the label array.
8. `MRC format version` : The `nversion` field should be 20140 or 20141 for compliance with the MRC2014 standard.
9. `Extended header type` : If an extended header is present, the `exttyp` field should be set to indicate the type of extended header.
10. `Data statistics` : The statistics in the header should be correct for the actual data in the file, or marked as undetermined.
11. `File size` : The size of the file on disk should match the expected size calculated from the MRC header.

During the checks, if the file is completely valid, the `mrc.validate` function returns `True`; otherwise, it returns `False`. Seriously invalid files trigger a `RuntimeWarning`. All data items listed above successfully passed the validation checks for the entire collection of cryo-EM density maps in the Cryo2StructData. This affirms the dataset’s full compliance with the MRC2014 format, confirming its validity as a set of density maps and masks suitable for application in AI-based methodologies.

### Validation using Deep Learning

To further validate the utility and quality of Cryo2StructData, we trained and test two deep transformer-based models^35^ on Cryo2StructData to predict backbone atoms and amino acid types from density maps respectively. Moreover, we used predicted C*α* atom positions and probabilities of 20 amino acid types to construct a Hidden Markov Model (HMM) model^35,36^ whose hidden states represent predicted C*α* atoms. The HMM model was used to align protein sequences with predicted C*α* positions to build the backbone structures of proteins via a customized Viterbi algorithm^35,37^.

#### Training and Testing Deep Transformer Models on Cryo2StructData

One transformer model was trained to classify each voxel of a sub-grid of a density map into one of four different classes representing three backbone atoms (C*α*, C and N) and absence of any backbone atoms. Another model was trained to classify each voxel of the sub-grid into one of twenty-one different amino acid classes representing twenty different amino acids and unknown or absence of amino acid. Additionally, we also trained a transformer model to predict the secondary structure of voxels, which was not used by build the HMM. In the inference stage, the predictions for all the sub-grids of a full density map are combined to generate a prediction for the entire map. We trained different sets of models with different parameters (measured in millions), dataset sizes, and extraction layers, as shown in Table 3

**Table 3.** Training on the Cryo2StructData with different deep learning models and data sizes. Refer to Table 2 for details on training and validation splits. “AHeads” refers to the number of attention heads for each transformer layer. “Models” refers to the trained and released model for amino acid type (amino), atom type (atom), and secondary structure type (ss) prediction in the Cryo2StructData Dataverse^30,31^.

|  | Data Size | Params (millions) | Transformers | AHeads | Extraction layers | Models |
| --- | --- | --- | --- | --- | --- | --- |
| Small Train | 1,867 | 35.60 | 4 | 6 | [1, 2, 3, 4] | amino, atom, ss |
| Full Train v1 | 7,392 | 92.28 | 12 | 12 | [3, 6, 9, 12] | amino, atom |
| Full Train v2 + Seq | 7,392 | 179.46 | 24 | 12 | [3, 6, 12, 24] | amino, atom |

For the `Small Train`, we used 25% of the Cryo2StructData (small subsample^29^), with 4 transformer layers and 6 attention heads. For `Full Train v1`, we utilized all the training and validation data from the Cryo2StructData (full data^30^), with 12 transformer layers and attention heads. We used the full trained models^30^ to construct HMMs (Section). The training and validation macro F1 score plots generated for `Full Train v1` and `Small Train` are shown in Figure 4. These plots illustrate the increase in F1 scores for both full and small training, providing further evidence that models trained on Cryo2StructData data can successfully learn and identify patterns in the data. ^1^

**Figure 4.**
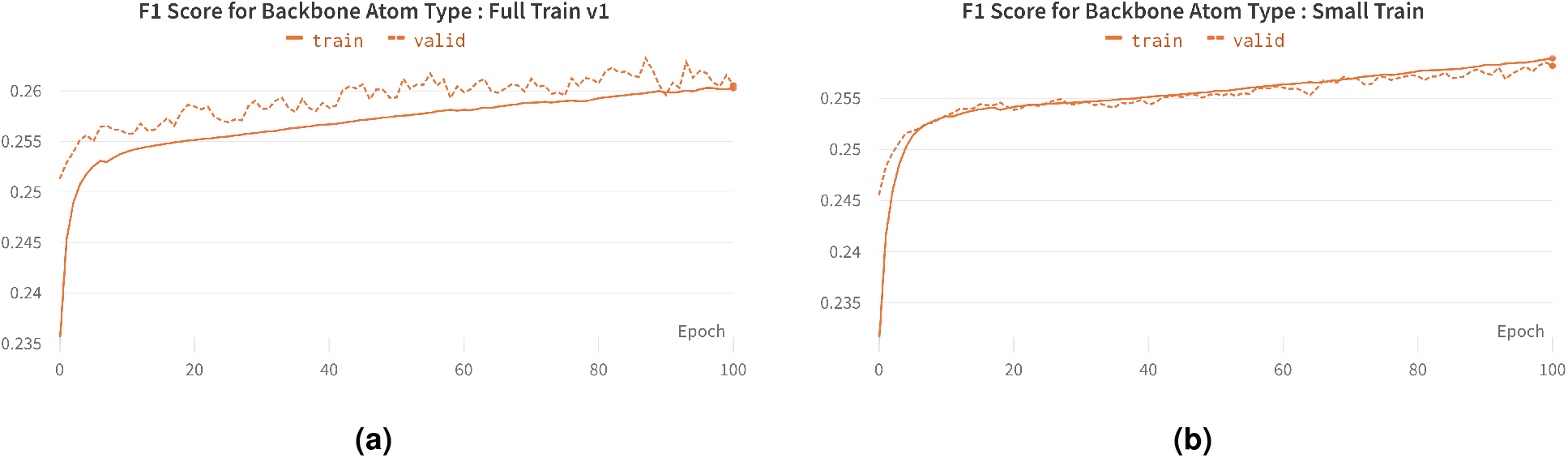
The training and validation F1 scores for transformer-model trained on Cryo2StructData. (a) Full Train v1. (b) Small Train.

For `Full Train v2 + Seq`, we utilized all the training and validation data from the Cryo2StructData (full data^30^), with 24 transformer layers and 12 attention heads. In addition, we integrated a protein language model (ESM-2^38^) generated sequence features into each protein density map, enhancing the sub-grid input with sequence information before passing it into the transformer layer. The per-residue representation is extracted using the ESM-2 pretrained model and then averaged to generate the per-sequence representation. We employed the ESM-2 pretrained model, which comprises 33 layers and 650M parameters, to generate embedding dimensions of 1,280. This computation was performed on the CPU of the Andes supercomputer^39^, utilizing its 256 GB RAM. To accommodate memory constraints, we utilized a sequence cutoff of 2,750 residues for each protein sequence. We extracted the encoded sequence representation from the 3^*rd*^, 6^*th*^, 12^*th*^, and the final 24^*th*^ layers. We noticed an increase in both validation recall and precision for the amino and atom type prediction tasks. A comparison of recall scores between `Full Train v1` and `Full Train v2 + Seq` is presented in Figure 5. By incorporating sequence information and enhancing the model’s complexity, the transformer-based model trained on the Cryo2StructData demonstrates improved learning. The sequence for each density map is made available within the Cryo2StructData, providing a convenient resource for AI-based researchers to utilize this complementary data and enhance the prediction capabilities of deep learning models.

**Figure 5.**
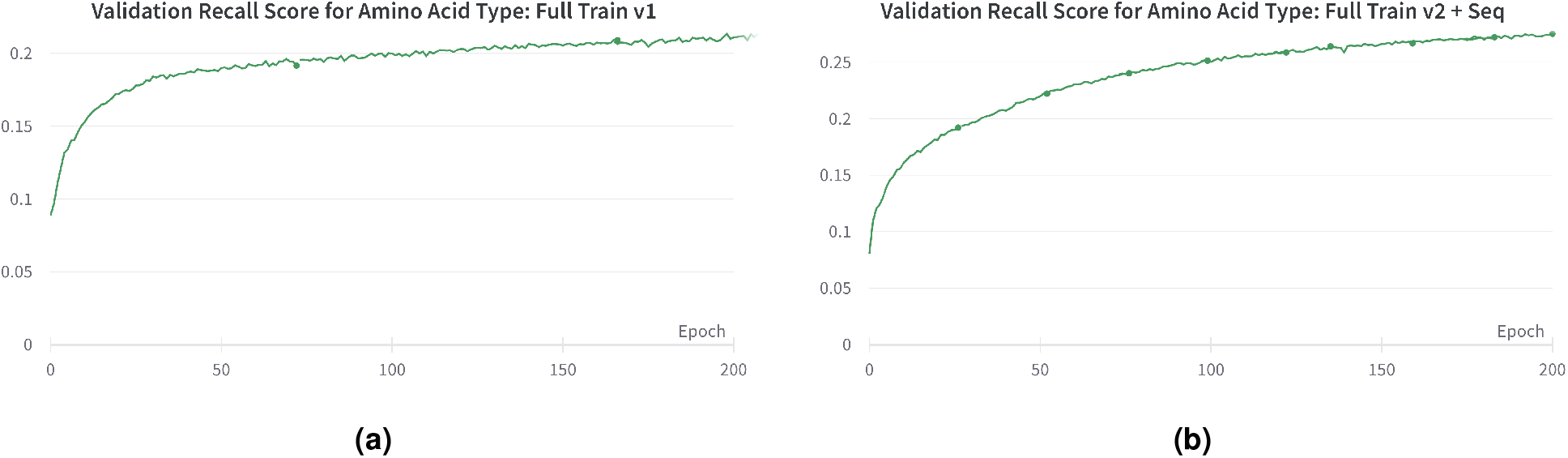
The validation recall scores for amino acid type prediction during training. (a) Full Train v1. (b) Full Train v2 + Seq. Incorporating sequence information to train the model improves the recall.

#### HMM-Guided Alignment of Cα Atoms and Protein Sequences

Connecting the predicted carbon-*α* atoms into protein chains and determining their amino acid types accurately is one of the most challenging aspects of building protein model from a cryo-EM density map. To address this challenge, we constructed a HMM^35^ from C*α* atoms and their amino acid types predicted by the deep transformer models. This model aligns the known protein sequences with the predicted hidden states denoting C*α* atoms in the HMM, allowing us to link C*α* atoms into protein chains and assign amino acid types to the atoms in a single step.

HMMs are generative models described by transition probability between hidden states and emission probabilities of generating observed symbols from hidden states. Specifically, the predicted C*α* atoms (voxels) from the atom type prediction are used as hidden states in the HMM, which are fully connected to generate the sequence of a protein. The amino acid emission probabilities of each C*α* hidden state are the normalized geometric mean of the probabilities of twenty different amino acid types predicted by the amino acid type prediction model and their background probabilities precomputed from the protein sequences in the training data. The geometric mean is computed as *a* × *b*, where *a* and *b* represent the predicted probability for an amino acid type and its background frequency, respectively. The transition probability between any two C*α* hidden states are calculated according to the euclidean distance (*x*) between two C*α* atoms based on the Gaussian distribution with a mean (*µ*) of 3.8047 and a standard deviation (*σ*) of 0.036 times a fine-tune able scaling factor (Λ) (Λ = 10).

Because it is not known which C*α* state generates the first amino acid of a protein chain, the HMM allows every C*α* hidden state to be the initial state. The probability for a C*α* state to be the initial state is equal to the probability of it emitting the first amino acid divided by the sum of the probability of every state emitting the first amino acid. The constructed HMM is then used by a customized Viterbi algorithm^35^ to compute the most likely path of aligning protein sequences with the C*α* hidden states, resulting a determined protein backbone structure. The customized Viterbi algorithm allows any C*α* state to occur at most once in the path because one C*α* position can be occupied by only one amino acid of a protein.

#### Evaluation Results on SARS COVID-19 Proteins

We tested the trained deep learning models on three SARS-CoV-2 proteins^40–42^ and one human p97 protein^43^. The voxel-wise predictions of C*α* atom were evaluated using F1-score (i.e., geometric mean of precision and recall of C*α* predictions). We used F1-score as it is a more balanced metric than the accuracy when there is a significant imbalance in the class distribution (i.e., the portion of voxels containing C*α* atoms is very small in this case). We compare the F1 scores of our predictions with those of the random predictions in Figure 6. The former is much better the latter.

**Figure 6.**
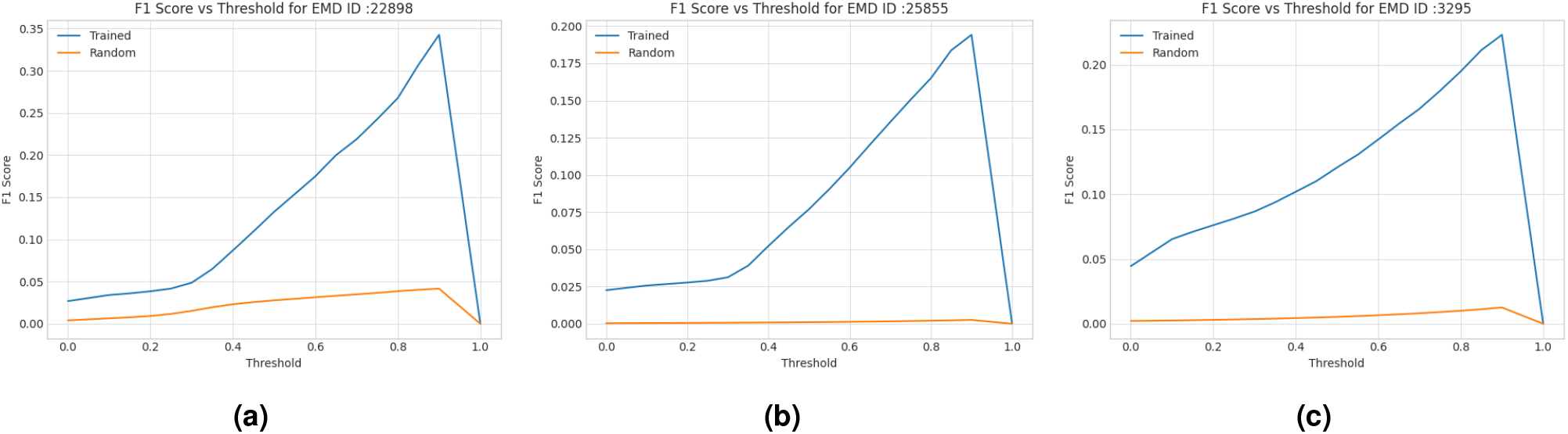
F1 scores of C*α* atom predictions for the density maps of two SARS COVID-19 (EMD-22898^40^, EMD-25855^42^) and a human p97 (EMD-3295^43^) test proteins. The curve in blue denotes how F1 score of the deep transformer trained on Cryo2StructData changes with respect to the threshold on predicted C*α* atom probabilities and the curve in orange the random predictions.

We further evaluated the C*α* backbones aligned by the HMM for the two COVID-19 proteins^40,42^ and the human p97 protein^43^ against the known protein structures as shown in Figures 8, 10, and 12. To adhere to the commonly practiced approach in the literature^18,27^, we used phenix.chain_comparison tool to compute the root mean squared distance (RMSD), the percentage of matching C*α* atoms, and the percentage of sequence identity. Phenix’s chain_comparison tool compares two structures to identify the number of matching C*α* atoms. Using this approach, it calculates the matching percentage, which represents the proportion of residues in the known structure that have corresponding residues in the reconstructed backbone structure. Similarly, it reports the sequence matching percentage indicating the percentage of matched residues that have the same amino acid type. The results in Table 4 shows that using the Cryo2StructData with the deep learning predictions and HMM alignment, we were able to determine the backbone structures of the four proteins with the good accuracy on average. Figures 7, 9, 11, 8, 10, and 12 shown in `Several examples of predicting C`*α* `atoms and reconstructing protein backbone structures` (Section) shows several good, detailed examples of predicting C*α* atoms and reconstructing protein backbone structures from density maps.

**Table 4.** Evaluation scores for predicted backbone structures for three SARS COVID-19 proteins^40–42^ and one human p97 protein (EMD-3295)^43^.

| EMDB | PDB | Residues | RMSD (↓) | Matching (% , ↑) | Sequence ID (% , ↑) |
| --- | --- | --- | --- | --- | --- |
| 22898 | 7KJR | 448 | 1.09 | 80.4 | 86.7 |
| 30210 | 7BV2 | 1036 | 1.31 | 69.7 | 73.0 |
| 25855 | 7TEY | 2703 | 1.42 | 71.6 | 85.0 |
| 3295 | 5FTJ | 4338 | 1.55 | 62.7 | 21.0 |
| Average |  |  | 1.34 | 71.1 | 66.42 |

**Figure 7.**
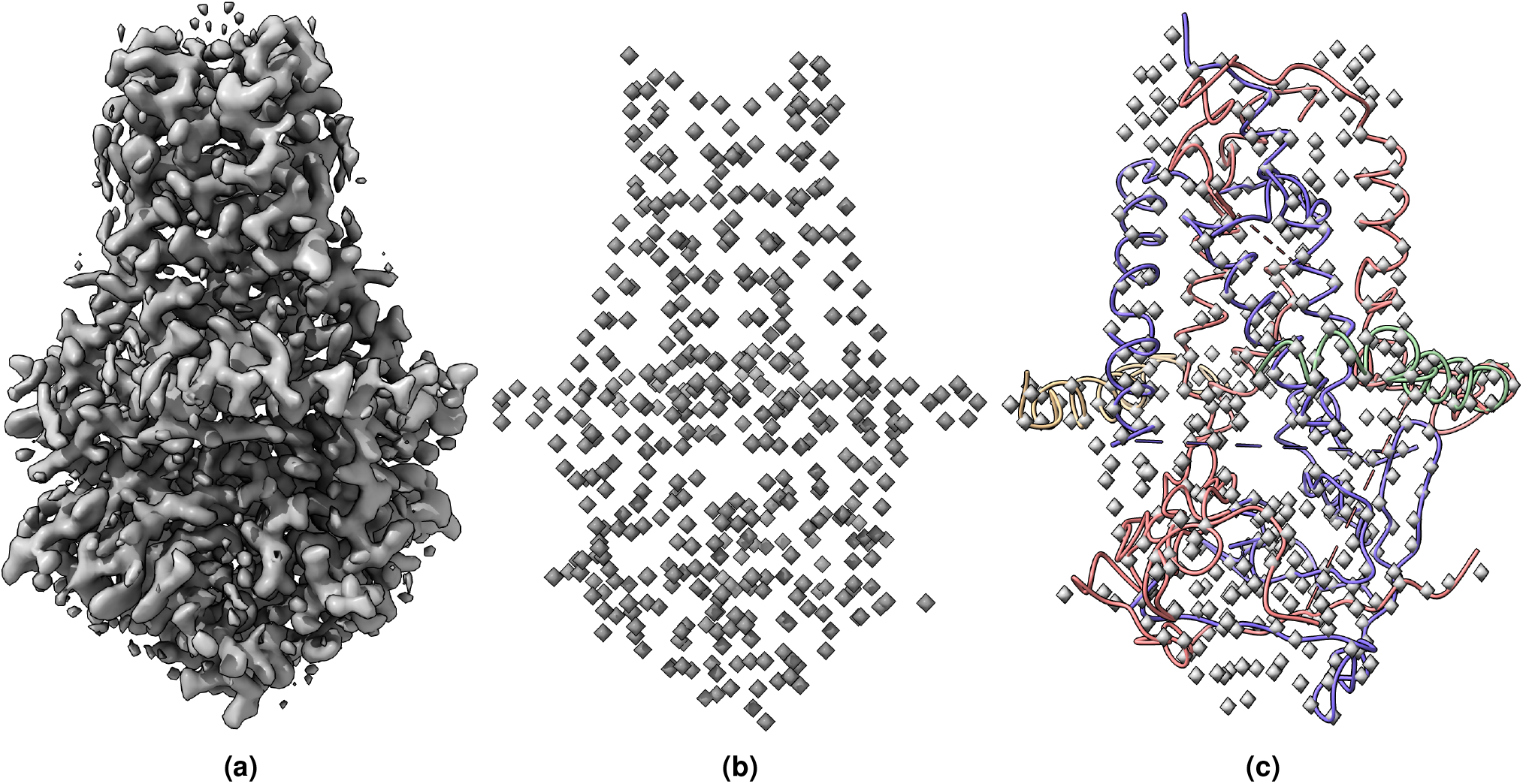
(a) The cryo-EM density map of SARS-CoV-2 ORF3a (EMD-22898) visualized at the recommended contour level of 0.7 (10.3 *σ*). The map dimension is 300 × 300 × 300 and has density values between −2.319 −3.909. The voxel dimensions is 0.727 × 0.727 × 0.727 (Å). (b) The true C*α* atom voxels (mask) extracted from the density map. (c) The predicted C*α* model is overlaid with true C*α* atom voxels.

**Figure 8.**
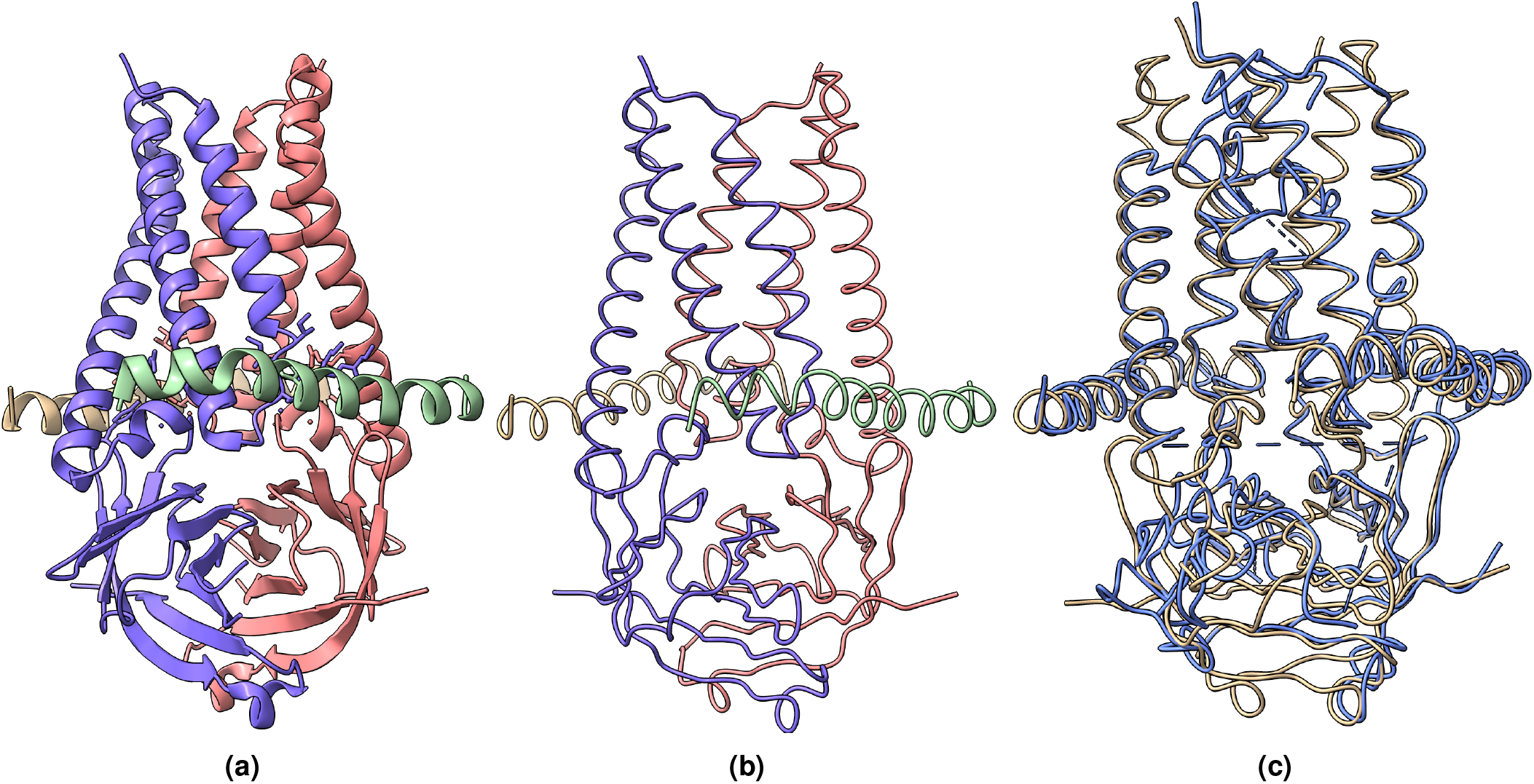
(a) The known protein structure corresponding to the density map EMD-22898 (PDB code 7KJR). The known structure has 448 residues. (b) The true C*α* backbone structure extracted from the known PDB structure. (c) The superimposition of the predicted backbone (blue) structure with the known backbone structure (gold).

**Figure 9.**
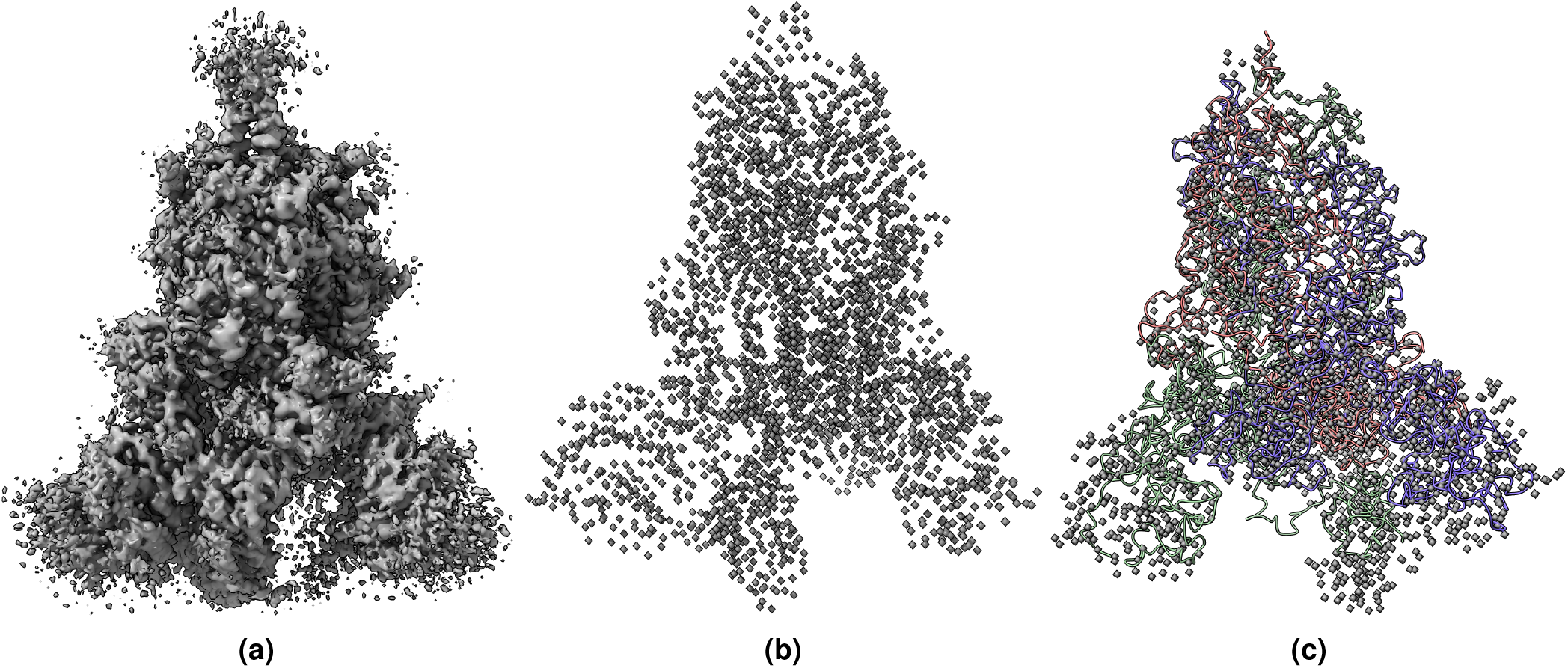
(a) The cryo-EM map of SARS-CoV-2 Delta (B.1.617.2) spike protein (EMD-25855) visualized at recommended contour level of 0.121 (5.0 *σ*). The map dimension is 400 × 400 v 400 and has density values between −0.467 −1.315. The voxel dimensions is 1 × 1 × 1 (Å). (b) The true C*α* atom voxels (mask) extracted from the density map. (c) The predicted C*α* model is overlaid with true C*α* atom voxels.

**Figure 10.**
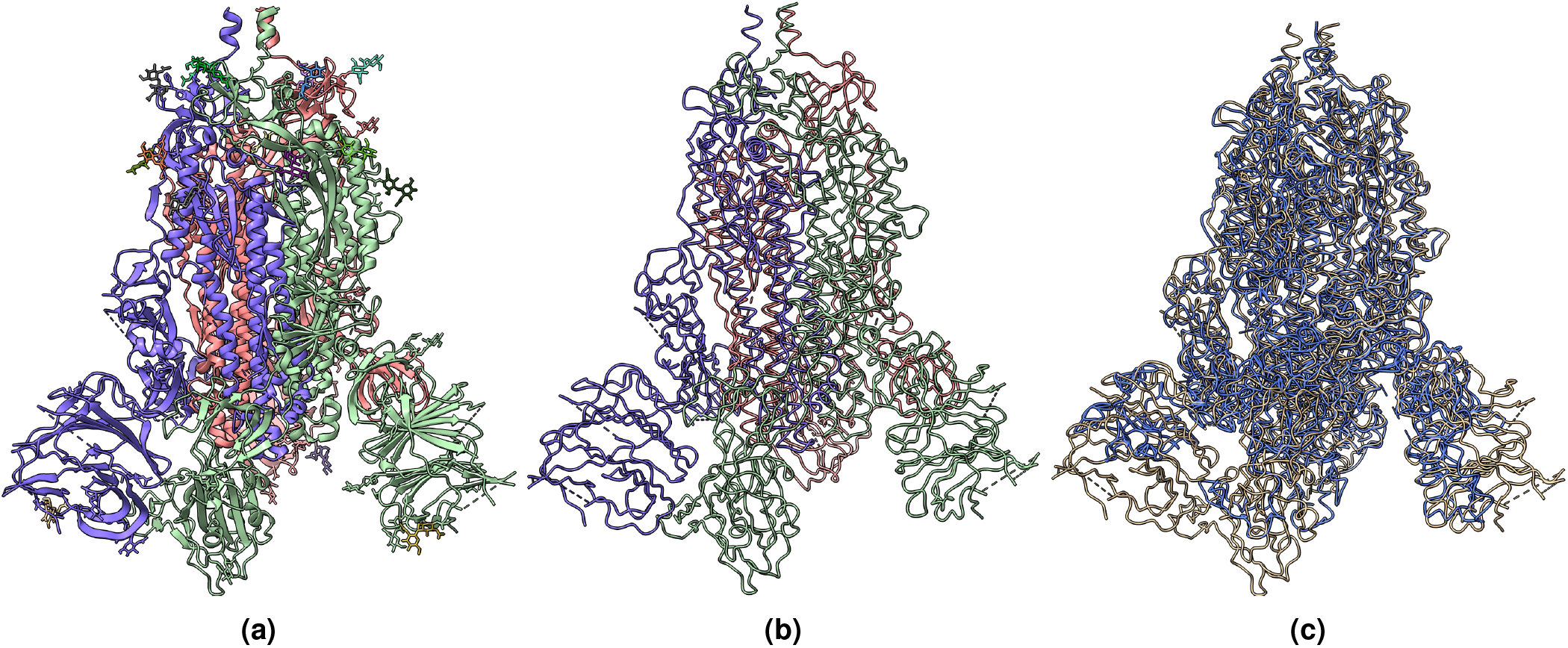
(a) The known protein structure of the density map EMD-25855 (PDB code 7TEY). The known structure has 2,703 residues. (b) The true C*α* backbone structure extracted from the known protein structure. (c) The superimposition of the predicted backbone structure (blue) with the known backbone structure (gold).

**Figure 11.**
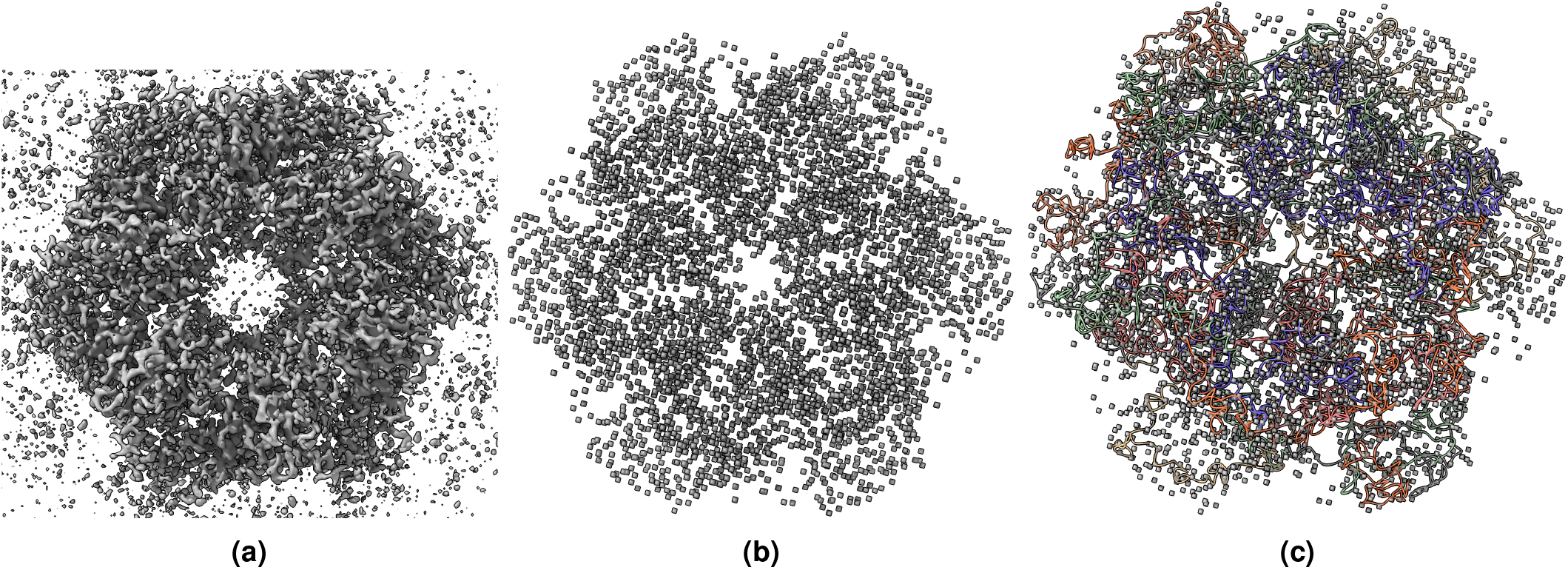
(a) The cryo-EM density map of human p97 bound to UPCDC30245 inhibitor (EMD-3295) visualized at recommended contour level of 0.0375 (3.4 *σ*). The map dimension is 312 × 312 × 312 and has density values between 0.085 − 0.118. The voxel dimensions is 0.637 × 0.637 × 0.637 (Å). (b) The true C*α* atom voxels (mask) extracted from the deposited density map. (c)The predicted C*α* model is overlaid with true C*α* atom voxels.

**Figure 12.**
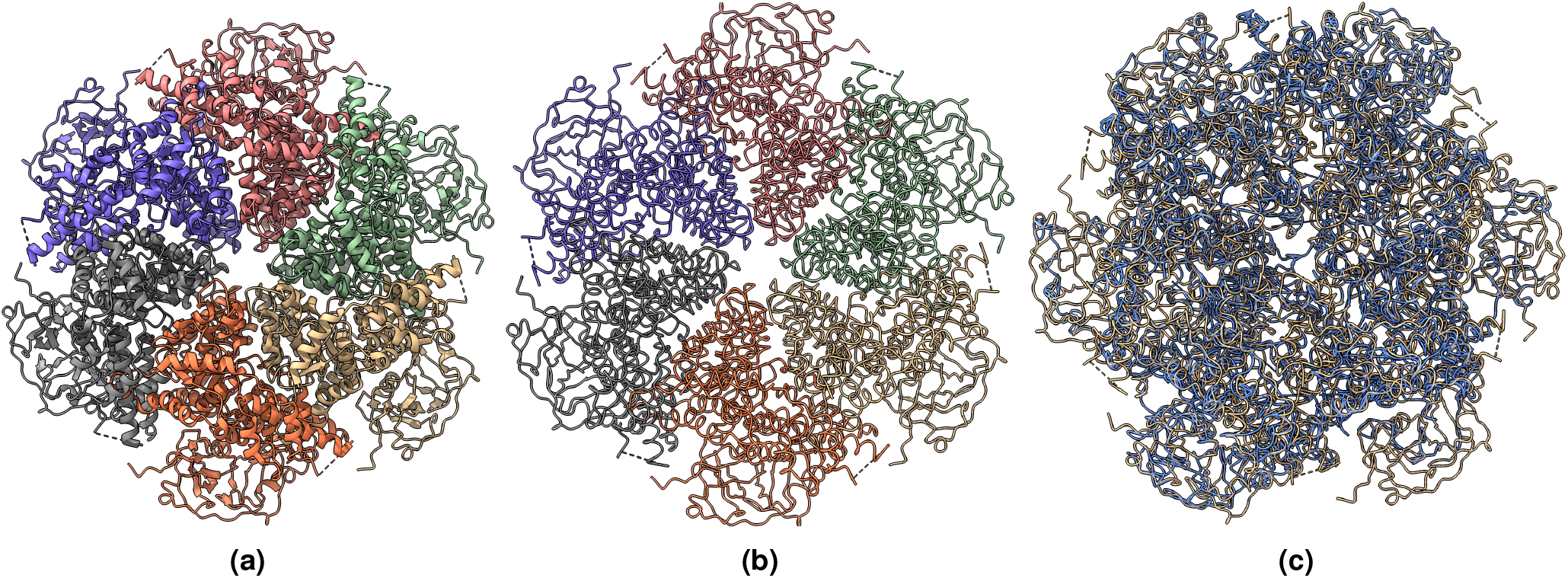
(a) The known protein structure of the density map EMD-3295 (PDB code 5FTJ). The structure has 4,338 residues. (b) The true C*α* atom backbone structure extracted from the known protein structure. (c) The superimposition of the predicted backbone structure (blue) with the known backbone structure (gold).

**Figure 13.**
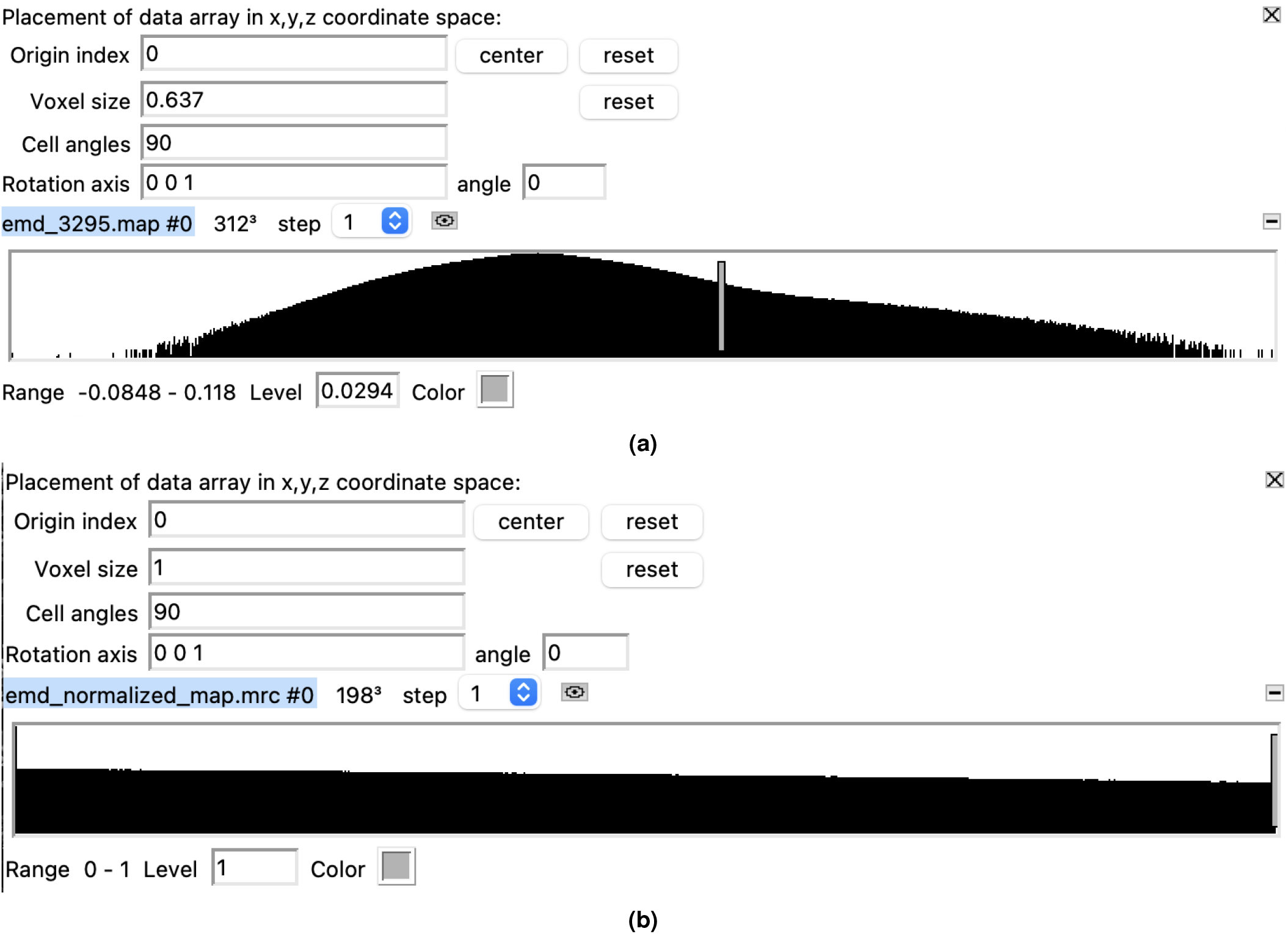
The cryo-EM density map of human p97 bound to UPCDC30245 inhibitor (EMD-3295). (a) Density values in raw cryo-EM density map. (b) Density values after preprocessing (resampled and normalized). The voxel size is 1 Å, and the density values are between [0 - 1]. The image is extracted from volume viewer of UCSF Chimera^25^.

## Usage Notes

Cryo2StructData is specifically designed for the task of building protein models from cryo-EM density maps. The prepared cryo-EM density maps available in Cryo2StructData Dataverse^12,29^ can be visualized using UCSF Chimera^25^. For the analysis of these maps, it is recommended to use the `mrcfile package`^26^ in Python.

The data preprocessing scripts in the Cryo2StructData GitHub repository can be modified or utilized to generate custom data tailored to user’s specific needs or reproduce the data generation process. The label generation process scripts are also included in the GitHub repository.

Cryo2StructData contains two types of *data splits*. The first type is resolution-based splits, and the second type is ID-based splits. We trained our deep learning model on the resolution-based splits dataset, as described in Table 2. Users have the option to use the splits we have created or can create their own splits based on their needs. Additionally, we provide a smaller downsampled version^29^ of the entire Cryo2StructData for users who do not have access to large computational resources. The split-based information is available in Cryo2StructData Dataverse^30,31^.

The provided *trained deep learning models*^30,31^ for atom and amino acid-type prediction, are essential for predicting voxels within the input cryo-EM density maps. The Cryo2StructData GitHub repository contains a deep learning inference program that utilizes these trained models for voxel-wise prediction. Our novel *Hidden Markov Model* as detailed in `HMM-Guided Alignment of C`*α* `Atoms and Protein Sequences` (Section), a fast and efficient method for aligning predicted voxels to form a protein backbone structure in a single step, is written in C++ and is available in the Cryo2StructData GitHub repository. Additionally, both the Cryo2StructData Dataverse and the Cryo2StructData GitHub repository contain detailed information about the dataset and instructions for building the protein backbone model from cryo-EM density map, enabling users to seamlessly utilize the data and code.

The integration of *ESM-2*^*38*^ *generated sequence features* into the protein density maps has yielded promising results, notably improving validation recall and precision for amino acid type prediction tasks, as shown in Figure 5. This plot demonstrates the impact of incorporating sequence information into the density map. The sequence for each cryo-EM density map is made available within the Cryo2StructData Dataverse^12,29^, providing a convenient resource for researchers to utilize this complementary data and enhance the prediction capabilities of deep learning models.

The cryo-EM density map data structure is prepared as a 3D (three dimension) grid of values organized into layers of rows and columns. The voxel grids can be inferred as a regularly spaced point cloud where each point is a voxel. Other approaches, such as modeling input cryo-EM density maps as 3D point clouds and leveraging 3D Graph Neural Networks^44^, can be considered.

The dataset of this scale allows researchers to train and test robust and powerful deep learning models to predict the positions of protein backbone atoms (e.g., C*α* atoms) and their amino acid types in 3D cryo-EM density maps, which can be linked together to build 3D protein models from scratch without using any known structural information as templates. The AI-powered de novo modeling of protein structures from cryo-EM density maps will significantly extend the capability and efficiency of cryo-EM techniques to solve the structures of large protein complexes and assemblies.

## Code availability

The source code and instructions to reproduce our results are freely available at https://github.com/BioinfoMachineLearning/cryo2struct. To keep the data files of Cryo2StructData permanent, we published all data to the Harvard Dataverse (https://dataverse.harvard.edu/dataverse/Cryo2StructData), an online data management and sharing platform with a permanent Digital Object Identifier number for each dataset. The Cryo2StructData Dataverse comprises the Full Cryo2StructData, referred to as `Cryo2StructData: Full Dataset`^12^, along with its associated trained deep transformer model and data splits, referred as `Cryo2StructData: Trained Model and Data Splits (Full)`^30^. Similarly, within the Cryo2StructData Dataverse, we find the Small Subsample of the complete Cryo2StructData, denoted as `Cryo2StructData: Small Subsample Dataset`^29^, accompanied by its respective trained deep transformer model and data splits, recognized as `Cryo2StructData: Trained` `Model and Data Splits (Small Subset)`^31^. Finally, the test dataset has been made available as `Cryo2StructData: Test Dataset`^28^.

## Acknowledgements

This work was supported in part by NIH grants (R01GM146340 and R01GM093123), Department of Energy grants (DE–SC0020400 and DE–SC0021303), and an NSF grant (DBI2308699). This research also used the computing resource of the Oak Ridge Leadership Computing Facility, which is a DOE Office of Science User Facility supported under Contract DE-AC05-00OR22725.

## Author contributions statement

J.C. conceived the project. J.C. and N.G. designed the methods. N.G. implemented and tested the methods. N.G. conducted the experiment and collected the data. N.G. and J.C. analyzed the data. N.G. and J.C. wrote the manuscript. L.W. revised the manuscript.

## Figures

### Several examples of predicting C*α* atoms and building protein backbone structures

Figures 7, 9, 11 visualize and analyze the predicted C*α* backbone structures of three proteins with respect to the true C*α* atoms of the density maps. The predicted backbone models were built by the deep transformer trained on Cryo2StructData and Hidden Markov Model. Figures 8, 10, 12 compare the modeled backbone structures of the same three proteins with their true backbone structures extracted from the known protein structures. We used UCSF ChimeraX^25^ for the visualization of the cryo-EM density maps and the protein structures. Generally, the modeled backbone structures match the true C*α* atoms in the density maps and the true backbone structures reasonably well.

## Footnotes

1 The trained models for both full and small versions are accessible in the Cryo2StructData Dataverse^30,31^, with the corresponding source code available on the Cryo2StructData GitHub repository.

## References

1. Boadu, F., Cao, H. & Cheng, J. Combining protein sequences and structures with transformers and equivariant graph neural networks to predict protein function. bioRxiv 2023–01 (2023).

2. Dhakal, A., McKay, C., Tanner, J. J. & Cheng, J. Artificial intelligence in the prediction of protein–ligand interactions: recent advances and future directions. Briefings Bioinforma. 23, bbab476 (2022).

3. Bai, X.-C., McMullan, G. & Scheres, S. H. How cryo-em is revolutionizing structural biology. Trends biochemical sciences 40, 49–57 (2015).

4. Kühlbrandt, W. The resolution revolution. Science 343, 1443–1444 (2014).

5. Iudin, A. et al. EMPIAR: the Electron Microscopy Public Image Archive. Nucleic Acids Res. 51, D1503–D1511, 10.1093/nar/gkac1062 (2022). https://academic.oup.com/nar/article-pdf/51/D1/D1503/49645431/gkac1062.pdf.

6. Dhakal, A., Gyawali, R., Wang, L. & Cheng, J. A large expert-curated cryo-em image dataset for machine learning protein particle picking. Sci. Data 10, 392 (2023).

7. Lawson, C. L. et al. Emdatabank unified data resource for 3dem. Nucleic acids research 44, D396–D403 (2016).

8. Jumper, J. et al. Highly accurate protein structure prediction with alphafold. Nature 596, 583–589 (2021).

9. Giri, N., Roy, R. S. & Cheng, J. Deep learning for reconstructing protein structures from cryo-em density maps: Recent advances and future directions. Curr. Opin. Struct. Biol. 79, 102536 (2023).

10. Giri, N. & Cheng, J. Improving protein–ligand interaction modeling with cryo-em data, templates, and deep learning in 2021 ligand model challenge. Biomolecules 13, 132 (2023).

11. Berman, H. M. et al. The Protein Data Bank. Nucleic Acids Res. 28, 235–242, 10.1093/nar/28.1.235 (2000). https://academic.oup.com/nar/article-pdf/28/1/235/9895144/280235.pdf.

12. Giri, N. & Cheng, J. Cryo2StructData : Full Dataset, 10.7910/DVN/FCDG0W (2023).

13. Si, D. et al. Deep learning to predict protein backbone structure from high-resolution cryo-em density maps. Sci. reports 10, 1–22 (2020).

14. Tang, G. et al. Eman2: an extensible image processing suite for electron microscopy. J. structural biology 157, 38–46 (2007).

15. Alnabati, E., Terashi, G. & Kihara, D. Protein structural modeling for electron microscopy maps using vesper and mainmast. Curr. Protoc. 2, e494 (2022).

16. Wriggers, W. Using situs for the integration of multi-resolution structures. Biophys. reviews 2, 21–27 (2010).

17. Cheng, Y., Grigorieff, N., Penczek, P. A. & Walz, T. A primer to single-particle cryo-electron microscopy. Cell 161, 438–449 (2015).

18. Pfab, J., Phan, N. M. & Si, D. Deeptracer for fast de novo cryo-em protein structure modeling and special studies on cov-related complexes. Proc. Natl. Acad. Sci. 118, e2017525118 (2021).

19. Jamali, K. et al. Automated model building and protein identification in cryo-em maps. bioRxiv 2023–05 (2023).

20. Mostosi, P., Schindelin, H., Kollmannsberger, P. & Thorn, A. Haruspex: a neural network for the automatic identification of oligonucleotides and protein secondary structure in cryo-electron microscopy maps. Angewandte Chemie Int. Ed. 59, 14788–14795 (2020).

21. He, J. & Huang, S.-Y. Emnuss: a deep learning framework for secondary structure annotation in cryo-em maps. Briefings bioinformatics 22, bbab156 (2021).

22. Zhang, X., Zhang, B., Freddolino, P. L. & Zhang, Y. Cr-i-tasser: assemble protein structures from cryo-em density maps using deep convolutional neural networks. Nat. methods 19, 195–204 (2022).

23. Maddhuri Venkata Subramaniya, S. R., Terashi, G. & Kihara, D. Protein secondary structure detection in intermediate-resolution cryo-em maps using deep learning. Nat. methods 16, 911–917 (2019).

24. Cheng, A. et al. Mrc2014: Extensions to the mrc format header for electron cryo-microscopy and tomography. J. structural biology 192, 146–150 (2015).

25. Pettersen, E. F. et al. Ucsf chimerax: Structure visualization for researchers, educators, and developers. Protein Sci. 30, 70–82 (2021).

26. Burnley, T., Palmer, C. M. & Winn, M. Recent developments in the CCP-EM software suite. Acta Crystallogr. Sect. D 73, 469–477, 10.1107/S2059798317007859 (2017).

27. Terwilliger, T. C., Adams, P. D., Afonine, P. V. & Sobolev, O. V. A fully automatic method yielding initial models from high-resolution cryo-electron microscopy maps. Nat. methods 15, 905–908 (2018).

28. Giri, N. & Cheng, J. Cryo2StructData : Test Dataset, 10.7910/DVN/2GSSC9 (2023).

29. Giri, N. & Cheng, J. Cryo2StructData : Small Subsample Dataset, 10.7910/DVN/CGUENL (2023).

30. Giri, N. & Cheng, J. Cryo2StructData : Trained Model and Data Splits (Full), 10.7910/DVN/SXNYRE (2023).

31. Giri, N. & Cheng, J. Cryo2StructData : Trained Model and Data Splits (Small Subset), 10.7910/DVN/DTV4JF (2023).

32. Pearson, W. R. & Lipman, D. J. Improved tools for biological sequence comparison. Proc. Natl. Acad. Sci. 85, 2444–2448 (1988).

33. Sievers, F. & Higgins, D. G. Clustal omega. Curr. protocols bioinformatics 48, 3–13 (2014).

34. Burnley, T., Palmer, C. M. & Winn, M. Recent developments in the CCP-EM software suite. Acta Crystallogr. Sect. D 73, 469–477, 10.1107/S2059798317007859 (2017).

35. Giri, N. & Cheng, J. De novo atomic protein structure modeling for cryo-em density maps using 3d transformer and hidden markov model. bioRxiv 0, 0 (2024).

36. Rabiner, L. & Juang, B. An introduction to hidden markov models. ieee assp magazine 3, 4–16 (1986).

37. Forney, G. D. The viterbi algorithm. Proc. IEEE 61, 268–278 (1973).

38. Lin, Z. et al. Language models of protein sequences at the scale of evolution enable accurate structure prediction. bioRxiv (2022).

39. Gao, M. et al. High-performance deep learning toolbox for genome-scale prediction of protein structure and function. In 2021 IEEE/ACM Workshop on Machine Learning in High Performance Computing Environments (MLHPC), 46–57 (IEEE, 2021).

40. Kern, D. M. et al. Cryo-em structure of sars-cov-2 orf3a in lipid nanodiscs. Nat. structural & molecular biology 28, 573–582 (2021).

41. Yin, W. et al. Structural basis for inhibition of the rna-dependent rna polymerase from sars-cov-2 by remdesivir. Science 368, 1499–1504, 10.1126/science.abc1560 (2020). https://www.science.org/doi/pdf/10.1126/science.abc1560.

42. Saville, J. W. et al. Structural and biochemical rationale for enhanced spike protein fitness in delta and kappa sars-cov-2 variants. Nat. communications 13, 742 (2022).

43. Banerjee, S. et al. 2.3 Å resolution cryo-em structure of human p97 and mechanism of allosteric inhibition. Science 351, 871–875, 10.1126/science.aad7974 (2016). https://www.science.org/doi/pdf/10.1126/science.aad7974.

44. Bronstein, M. M., Bruna, J., Cohen, T. & Veličković, P. Geometric deep learning: Grids, groups, graphs, geodesics, and gauges. arXiv preprint arXiv:2104.13478 (2021).

